# Structure of transmembrane helix 8 and possible membrane defects in CFTR

**DOI:** 10.1101/218792

**Authors:** Valentina Corradi, Ruo-Xu Gu, Paola Vergani, D. Peter Tieleman

## Abstract

The cystic fibrosis transmembrane conductance regulator (CFTR) is an ion channel that regulates the flow of anions across epithelia. Mutations in *CFTR* cause cystic fibrosis. CFTR belongs to the ATP-Binding Cassette (ABC) transporter superfamily, and gating is controlled by phosphorylation and ATP binding and hydrolysis. Recent ATP-free and ATP-bound structures of zebrafish CFTR revealed an unwound segment of transmembrane helix (TM) 8, which appears to be a unique feature of CFTR not present in other ABC transporter structures. Here, by means of 1 μs long molecular dynamics simulations, we investigate the interactions formed by this TM8 segment with nearby helices, in both ATP-free and ATP-bound states. We highlight the structural role of TM8 in maintaining the functional architecture of the pore and we describe a distinct membrane defect that is found near TM8 only in the ATP-free state. The results of the MD simulations are discussed in the context of the gating mechanism of CFTR.

---

The cystic fibrosis transmembrane conductance regulator (CFTR) regulates anion flow through the apical membrane of epithelia (1). Defects in CFTR cause cystic fibrosis, a disease in which the movement of ions and water across epithelia is compromised, allowing for thick secretions to accumulate mainly in the airways, digestive and reproductive systems (2). CFTR, although a member of the ATP-Binding Cassette (ABC) transporter superfamily, functions as a channel, switching between closed and open states (3).

An interesting feature of several recent CFTR structures is a region of transmembrane helix (TM) 8 that is partially unwound (residues 929-938 in zebrafish CFTR, zCFTR, corresponding to residues 921-930 in human CFTR, hCFTR), thus breaking the α-helical structure (4-6). This feature, common to the ATP-free and ATP-bound states, might underlie CFTR’s unique channel function (6). In order to investigate the interactions between this segment and nearby helices and how this segment can be partially unwound despite being in the membrane, we embedded the structure of zCFTR in the ATP-free and ATP-bound state in a POPC lipid bilayer and performed atomistic, 1 μs long, molecular dynamics (MD) simulations (Fig. 1).

**Figure 1.**
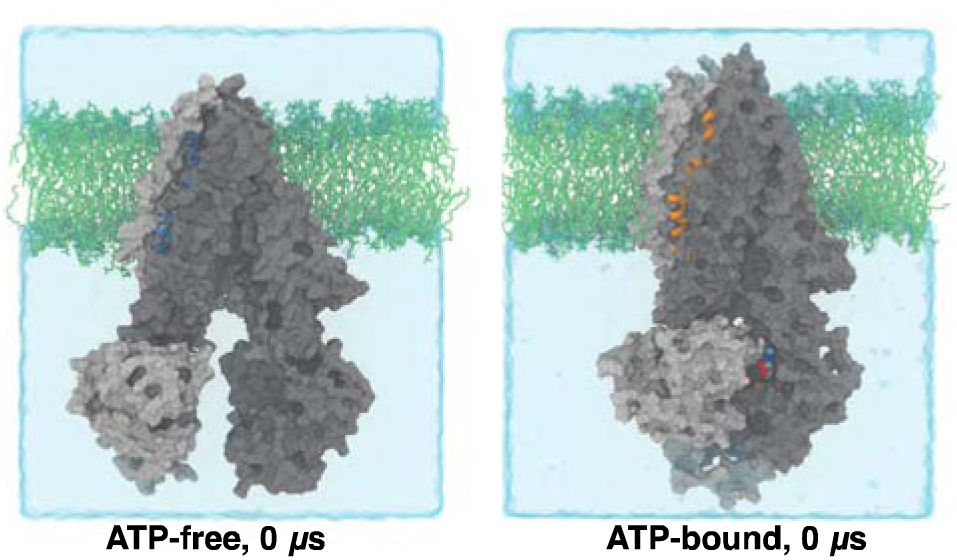
Simulation setup. The ATP-free (left) and ATP-bound zCFTR structures were embedded in a POPC bilayer (light green sticks). TMD1-NBD1 and TMD2-NBD2 are shown in dark and light surfaces, respectively. TM8 is highlightes in blue and orange cartoons, for the two systems. Water (light cyan surface) and lipids are clipped for clarity.

The conformation of individual domains (transmembrane domains, TMDs, and nucleotide binding domains, NBDs), based on root mean square deviation, inter-domain distances and secondary structure analyses, remains stable over 1 μs long MD simulations. In the ATP-free system we noticed a closing motion of the NBDs as also reported by others (7) (Fig. S1-S4).

The unwound segment of TM8 (Fig. S5) remains stable during the simulations in both systems (Fig. S6). The extracellular end of TM8, due to its different orientation between the ATP-free and the ATP-bound state (6), establishes interactions with residues of TM6 only in the presence of ATP (Y925 - S342 (TM6)) (Fig. 2A, S7, Table 1). Further along TM8, we identify several residues that interact with nearby helices in both states, including the E932 - R348 (TM6), S933 - F312 (TM5), L935 - Q1004, and Y305 - Q1004 (TM9) pairs (Fig. S7, Table 1). After the helix break, R941 establishes hydrogen bonds with E871 (TM7) only in the ATP-bound system (Fig. S7, Table 1).

**Figure 2.**
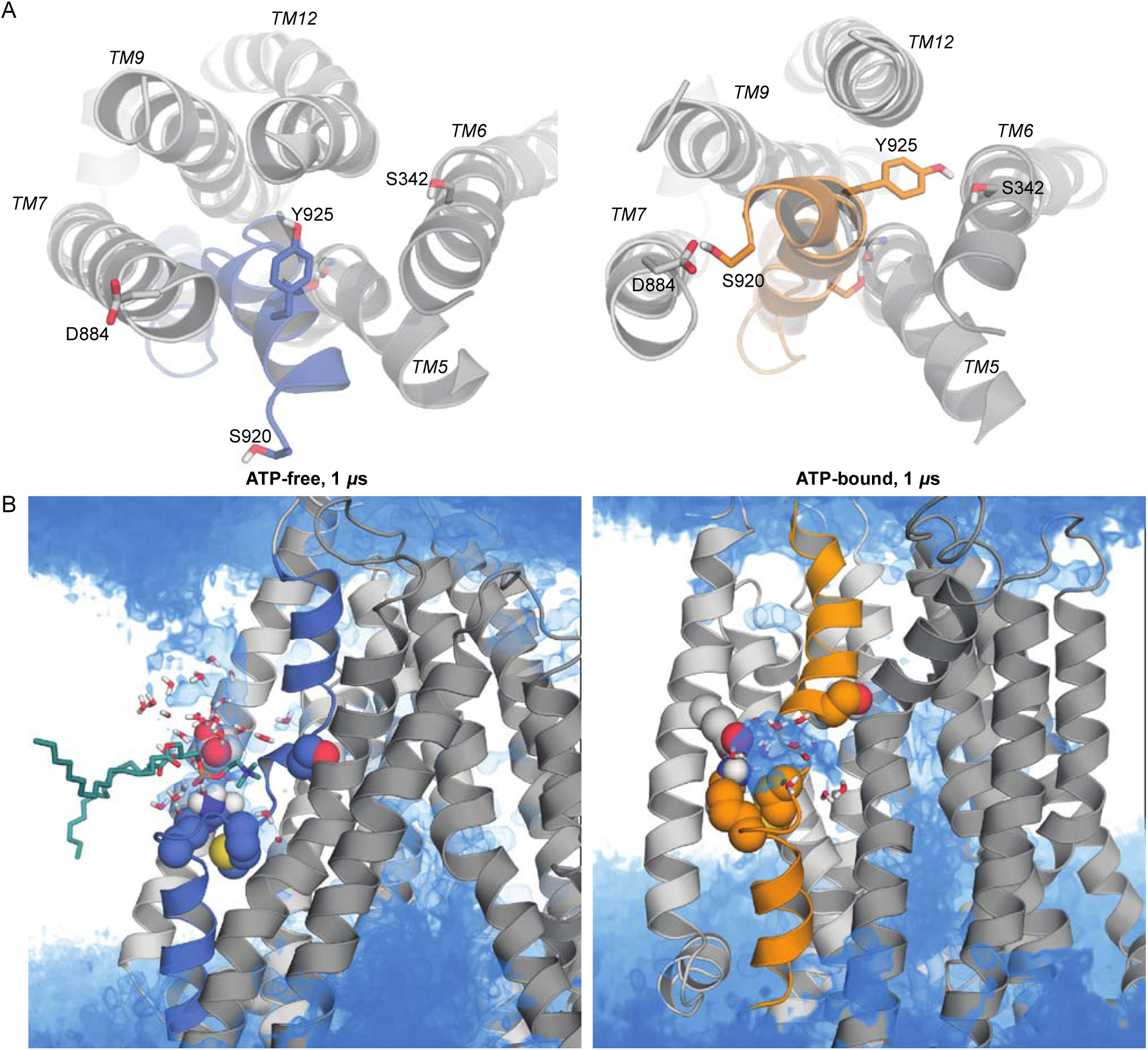
TM8 orientation and membrane defect. A. Extracellular view of TM8 and neighbouring helices in the ATP-free and ATP-bound system, after 1 μs of simulation time. B. The number density of water molecules, calculated during the last 50 ns of the simulation, is shown as a blue volume map and overlapped to the last frame of the simulation. TMD1 and TMD2 are shown in dark and light gray cartoons, respectively, with TM8 highlighted in blue (ATP-free system) and orange (ATP-bound system) cartoons. The side chains of E871, R941, M937 and S933 are shown as spheres, and the nearby water molecules within a 8 Å cut-off from R941 and E871 are shown as sticks. A lipid molecule with the head group interacting with R941 is shown for the ATP-free system (B) in dark-teal sticks.

**Table 1.**
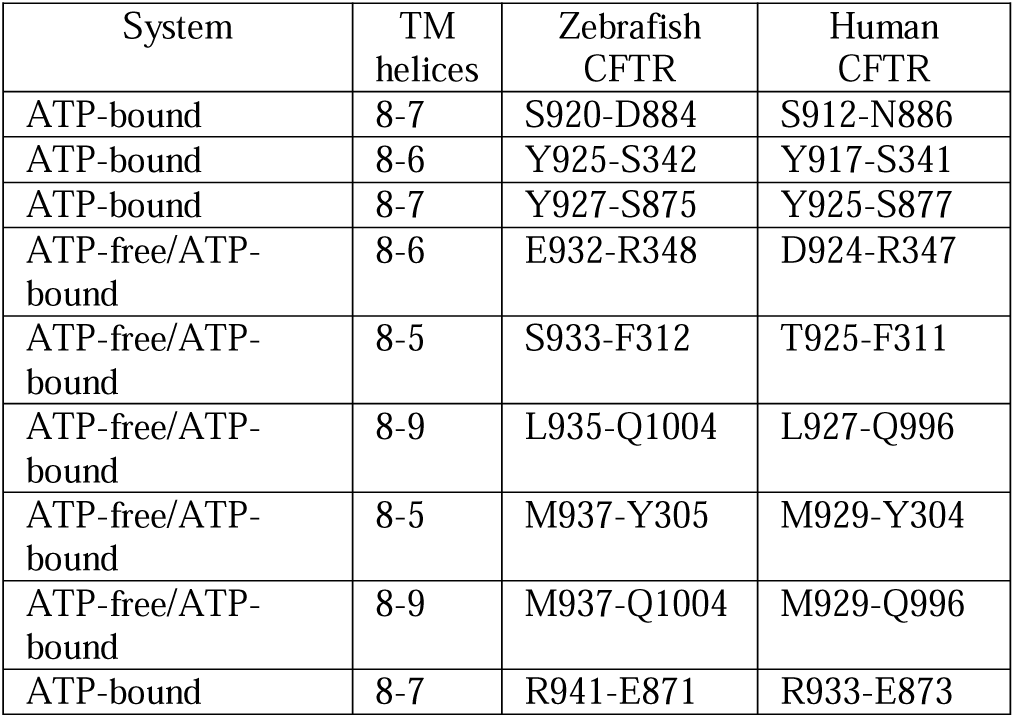
Summary of the interactions between TM8 and nearby helices found in the ATP-free and ATP-bound systems. The human CFTR numbering is given as a reference.

The unfolded segment of TM8 exposes the backbone and side chains of residues 930-941 to the hydrophobic region of the lipid bilayer, leading to the penetration of water and lipid headgroups inside the membrane (Fig. S8-S9). The number of water molecules found within a cut-off radius of 8 Å from R941 and E871 is doubled in the ATP-free system and accompanied by up to 5 lipid phosphate groups (Figure S8-S9). The phosphate atoms of the lipids take ca. 100 ns to reach the bilayer middle to start interacting with the side chain of R941 (Fig. S8). As a consequence, a clear water channel forms between the extracellular bulk solution and the unwound TM8 segment (Fig. 2B). Water molecules from the main cavity of the channel can also contribute to the pool of water molecules. In the ATP-bound system water channels from the extracellular side to the unwound segment of TM8 are not observed, while water molecules from the inner cavity of CFTR can still reach this region of TM8 (Fig. 2B).

In all ABC exporter structures the TMs are organized in a roughly symmetrical pattern, with no breaks in their helical structure (8). The unwound segment of TM8 is then a unique feature of CFTR (4-6) that suggests a solution for a long-standing puzzle: Why are the transmembrane helices known to line the permeation pathway not symmetrically distributed between TMD1 and TMD2? In particular, cysteine accessibility studies performed on TM7 and TM1 gave very different results (9-11), indicating that the latter but not the former, formed part of the permeation pathway (12,13). Chen and colleagues discovered that the unwound portion of TM8 and the resulting bend position the extracellular end of TM8 closer to the permeation pathway (4-6). This displaces TM7 further from the protein’s central axis, thus explaining the lack of contribution of TM7 to the anion permeation pathway.

Contacts between TM8 and TM6 are known to be required for maintaining the functional architecture of the channel, as in the case of residues E932 and R348 (14,15), whose interaction is retrieved in both ATP-bound and ATP-free simulation systems. TM8 also interacts with TM5, intracellular to the kink at F312, allowing dynamic interactions that stabilize the unwound segment of TM8 in both simulation systems. Overall, although not many of the residues that form molecular contacts with the unwound stretch of TM8 have been studied functionally, the network of molecular contacts between TM8 and the nearby helices investigated in our simulations can explain the stability of the unwound segment in both the ATP-free and the ATP-bound state, and is consistent with an active role of TM8 in the functional structure of the channel.

However, the current ATP-bound structure is not a fully-open channel, and the protrusion of TM8 residues towards the central pore interferes with anion flow, confirmed by the fact that an interaction is formed between TM8 and TM6, located in the narrowest portion of the pore (3,16). Thus, within the gating cycle of CFTR, the ATP-dependent interaction between the N-terminal of TM8 and TM6 might be transient, as part of an intermediate state that provides an additional gate preventing anion flow through the channel. While ATP-binding and NBD dimerization is coupled with the channel entering the “open burst” state (17), “flicker” closures of the pore occur within the open burst, with timescales that are not consistent with ATP hydrolysis (18-20). As suggested by Zhang and colleagues (6), the ATP-bound cryo-EM structure might have captured a flicker-closed state, artefactually stabilized by experimental conditions. The alternative hypothesis proposed by these authors, that the extracellular TM8 (and TM12) might reorientate within microseconds to a position allowing fast anion flow seems unlikely, given the relatively stable position of these secondary structure elements and the persistence of the Y925-S342 interaction in our simulations (Figures S3-S4, S6, S7).

The ATP-free, inward-facing structure shows an unusual lipid organization near TM8. Here, a strong membrane defect forms near R941, and E871, while no lipid head groups were found in proximity of these residues for the ATP-bound system. Although the presence of charged residues buried along TM helices is not common, there are examples where such residues play important biological roles, for instance in integrin receptors (21). The presence of a membrane defect with lipid headgroups and water molecules located in the bilayer middle is associated with a substantial energetic cost, although this cost can be compensated by a sufficiently hydrophobic sequence in the transmembrane helix (22-24). In the ATP-bound system, R941 and E871 engage in stable salt-bridge interactions that can screen the charges from the hydrophobic environment of the lipid tails, requiring no lipids to be pulled in to interact with R941. In the ATP-free system, on the other hand, this screening effect does not take place. Thus, the membrane defect and the cone of water molecules compensate for the charges of R941 and E871. This defect reaches the bilayer middle, and creates a side channel that occasionally allows the exchange of water molecules between the extracellular environment and the inner cavity of CFTR. Because the altered relative orientation of TM8 closes the permeation pathway in the ATP-bound cryo-EM structure (6), it is possible that, for the pathway to really open, the membrane defect might need to be restored, so as to support a TM8 orientation similar to that seen in the ATP-free structure. Further functional studies are however required.

Combined, our simulations reveal the stability of the unwound segment of TM8 in the ATP-bound and ATP-free state of CFTR. The simulations also highlight a pool of interactions, common to both states, between TM8 and nearby helices, that prevent further folding or unfolding of TM8 and contribute to the architecture of the channel. A strong membrane defect near R941 and E871 characterizes only the ATP-free system, but a similar membrane environment might provide structural support to a reorientation of TM8, opening the permeation pathway in the ATP-bound system too. It is interesting to note that such a reorientation would effectively shield a large portion of TM5 from the permeation pathway, in both closed and open states, as experimentally observed (9,13). However, we should also consider the possibility of structural artefacts, due to the lack of a native lipid environment or low temperature, affecting the conformation adopted by CFTR in the cryo-EM structures. While at present we can only speculate on whether the membrane disruption could be a feature of a fully open state, or perhaps could provide an interaction site for CFTR-interacting proteins, this study supports the development of experimental hypotheses to evaluate the contribution of TM8 residues to the gating mechanism of CFTR.

## SUPPORTING MATERIAL

Methods and Supporting Materials are available at

## AUTHOR CONTRIBUTIONS

V.C., P.V., D.P.T. designed the project; R.-X.G. perfomed the simulations and carried out the DSSP analysis; V.C. performed the remaining analyses and wrote the manuscript with P.V. and D.P.T, with contributions from R.-X.G.

## AKNOWLEDGEMENTS

This work was supported by the Canadian Institutes of Health Research. Additional support came from Alberta Innovates Health Solutions (AIHS) and Alberta Innovates Technology Futures (AITF). R.X.G. is supported by fellowships from AIHS and the Canadian Institutes for Health Research (funding reference number: MFE-140949). DPT holds the AITF Strategic Chair in (Bio)Molecular Simulation. Simulations were run on Compute Canada machines, supported by the Canada Foundation for Innovation and partners. This work was undertaken, in part, thanks to funding from the Canada Research Chairs program. PV thanks the Cystic Fibrosis Trust for support.

